# Unveiling Stimulus Transduction Artifacts in Auditory Steady-State Response Experiments: Characterization, Risks, and Mitigation Strategies

**DOI:** 10.1101/2024.05.30.596610

**Authors:** Jan Strobl, Vojtech Viktorin, Marek Piorecky, Inga Griskova-Bulanova, Jan Hubeny, Martin Brunovsky, Tomas Palenicek, Vlastimil Koudelka

## Abstract

This scientific paper addresses the potential risk of spurious responses in neuroscientific auditory steady-state response (ASSR) experiments attributed to transduction artifacts. The focus is particularly on click train stimuli, given their spectral content in the frequency range of interest (e.g., 40 Hz). Building upon a pilot experiment demonstrating the existence of the artifact in a phantom head, this study focuses on the characterization of stimulus artifacts in extended measurements with phantoms and the evaluation of associated risks in experiments involving human subjects. The investigation is divided into two parts: the first part scrutinizes stimulus artifact properties crucial for mitigation, while the second part assesses risks in ASSR experiments with human subjects based on the characterized artifact. The discussion covers stimulus characterization, experimental setups with phantoms, and experiments with human subjects, exploring potential sources of the artifact, its spatial properties, and the influence of re-referencing. The results reveal the role of headphone cables as a source of stimulus artifacts, along with the surprising impact of headphone transducers. The study emphasizes the need for careful experimental design and data analysis to avoid misinterpretations of stimulus artifacts as genuine brain responses in ASSR experiments.

## 1 Introduction

Auditory steady-state response (ASSR) is a response of the brain to a periodic auditory stimulation that is recorded with electroencephalography (EEG) or magnetoencephalography (MEG), and it is strongest at around 40 Hz [1]. These 40 Hz auditory steady-state responses reflect both the superposition of the middle-latency responses [2, 3] and a periodic resonant activity within the networks activated by the auditory periodic stimulation [4, 5]. The response originates within the auditory cortex and subcortical regions [6], with some contribution from a widely distributed network of sources within the frontal, motor, parietal, and occipital lobes [7, 8, 9]. At the cellular level, ASSRs are generated by the synchronous recruitment of inhibitory fast-spiking parvalbumin interneurons that are gamma-aminobutyric acid (GABA)-ergic cells receiving N-methyl-D-aspartate (NMDA)-dependent excitatory input from pyramidal cells [10, 11].

One of the main areas of research is the function of ASSRs as biomarkers for psychotic states, especially in schizophrenia [12]. The responses are diminished in phase synchronization and magnitude in patients with schizophrenia [13], bipolar disorder [14, 15, 16, 17], and also in high-risk subjects for psychotic onset [18].

Different types of stimulation may be used to elicit ASSR. The most commonly used stimulation is the “click train,” which is well-tested (e.g., [19, 20]). This type of stimulation results in pronounced and reliable responses [21, 22, 12] but may be unpleasant for the participant [23]. Another type of frequently employed stimulation is amplitude modulated (AM) tones [4].

When analyzing ASSRs, the primary focus is on the main response frequency, and the majority of studies do not focus on the higher harmonics of the ASSR. However, several attempts were made to evaluate higher order harmonics in response to 20 Hz stimulation, where response at 40 Hz is clearly visible [24]. Also, [25] assessed ASSRs to four click train stimulation frequencies (20, 30, 40, and 80 Hz). The inter-trial phase coherence (ITPC) of the time-frequency response is visible in the higher harmonic frequency of the response to 20 Hz click train stimuli in their study. This activity is comparable with the response at the fundamental frequency. Conversely, the same study did not detect higher harmonics in response to 30 Hz click train stimuli. Unfortunately, most of the previous studies also evaluate data within a frequency range of up to approximately 50 Hz only. Therefore, it is unclear whether a higher harmonic frequency response to 40 Hz stimuli is present. A higher harmonics response to 40 Hz click train stimuli was evident in studies by [9] and [26]. However, the use of AM stimulation [27] did not demonstrate the presence of higher harmonics in response to 40 Hz stimuli. McFadden et al. [22] reported that the higher harmonic frequency response to 40 Hz click train stimulation is stable across measurements, in contrast to responses to AM stimuli.

ASSRs are frequently obtained using EEG, which is very sensitive to surrounding interference (artifacts -signals not coming from the subject’s brain) [28]. Due to the use of headphones or earphones, ASSR experiments may be subjected to specific stimulus transduction artifacts (hereinafter referred to as - the stimulus artifact) in addition to common EEG artifacts. The source of the stimulus artifact is the headphone cable or headphone transducer [29]. Akhoun et al. [30] identified the headphone transducer as the worst source of stimulus artifact that is caused by the changes in the magnetic field arising from moving coils [31]. Alternatively, Campbell et al. [29] claim that a headphone cable is the primary source of this artifact as it conducts the acoustic stimulus like an electrical signal that may be registered in the immediate surroundings of the headphone cable. Therefore, the stimulus artifact is present only during the time of the ongoing stimulation and has the same frequency characteristics as the auditory stimulus itself[29, 30]. The power of this artifact increases with the increasing intensity of the stimulus. It also depends on the distance of the headphone transducer from the EEG electrodes, the orientation of the headphones cable and transducer with respect to orientation of the EEG wires and the hdEEG net, and the type of the headphone transducer [32, 33]. Akhoun et al. [30] reported that the EEG electrode impedance influenced the stimulus artifact in their study. It was not present in an artificial situation with zero impedance. On the other hand, the artifact was comparable for cases with normal (approximately 1000 Ω) and high (over 400 kΩ) impedance, suggesting that the influence of impedance is not profound.

It is crucial to disentangle the true ASSR effect originating in the brain from the transduction effect if present. However, it is difficult or even impossible to control the contribution of the transduction artifact post hoc.

Several methods were proposed for stimulus artifact suppression during the experiment. Prevention by suitable experimental design is one way to suppress the stimulus artifact. The increase in the distance between the headphone transducer and the EEG system is one of such experimental designs. Classic earphones with a plastic tube are used in some studies (for example, [34, 35]) to increase this distance. However, plastic tubes lead to acoustic dispersion, which may be a problem with click stimulus [29]. Using headphones with a different type of transducer may be a solution, but these headphones tend to be expensive [31].

Another option is to stimulate via speakers that do not create stimulus artifacts; however, acoustic wave reverberation may arise in a measuring cabin that is not sound-attenuated [36]. It should be noted that only a few studies employing ASSRs in research settings used speakers, and often those studies utilized unique protocols (for example, [37] compared the effect of ASSR in humans and monkeys and [38] applied transcranial alternating current stimulation during their ASSR experiment). Finally, a relatively effective way for stimulus artifact suppression is the shielding of the headphone cable, especially combined with grounding to a Faraday cage [29, 30, 39].

The stimulus artifact may also be suppressed to a certain level by correct data preprocessing. These methods are applied mainly in experiments utilizing AM stimuli. Some studies explain the stimulus artifact problem in these experiments by aliasing (see [33]). Averaging the brain responses to stimuli with opposite polarity is widely used for artifact suppression in studies using AM stimuli (for example, [40, 41]). The brain response creates envelopes the stimulus signal, which remains preserved [29, 42]. Unfortunately, this method cannot be adapted to studies with click train stimuli that contain white noise. The re-referencing method is applicable for suppression of the stimulus artifact originated by click train stimuli, i.e., the EEG channels are re-referenced to the electrode containing stimulus artifact and not con-taining a brain response [43, 44]. However, this requires extra hardware, which is a disadvantage of this method.

Several studies used more advanced techniques for stimulus artifact suppression. For example, [45] used earphones with plastic tubes for stimulation and, during data preprocessing, averaged the stimuli with opposite polarity. Campbell et al. [29] compared methods averaging the opposite phase stimuli, re-referencing EEG electrodes, and the headphone cable shielding. The authors recommended a combination of all compared methods and identified headphone cable shielding as the most effective means. Akhoun et al. [30] also recommended the headphone cable shielding with grounding to the Faraday cage. Brooke et al. [32] got similar results. In this study, the authors also described the significant influence of the change in transducer orientation on stimulus artifacts.

There is a particular reason the above-mentioned techniques may fail in the case of the click train stimuli. Firstly, the auditory entertainment, which is the main interest in the ASSR experiments, is phase-locked to the stimulus itself, i.e., to the 40 Hz stimulation train. Thereby, changing the polarity of the stimulus also changes the polarity of the brain response, and the averaging technique suppresses both the artifact and the true brain response. Secondly, the click train stimulus contains sub-harmonical spectral components, which are identical to the frequencies of interest, i.e. 40, 80, and 120 Hz.

Despite several existing studies on stimulus artifacts, most of them do not focus on the context of neuroscientific research, and the ASSR paradigm is rarely utilized in these works. Hence, the main aim of this study is to systematically investigate the possible occurrence of stimulus artifact in ASSR during click-induced stimulation, describing its presence, risks, and properties following up on our original pilot study [46], which confirmed the presence of this artifact in a phantom experiment.

We elaborated two experimental approaches to reach the aim. The first experimental design includes several measurements with a human head phantom, aiming at confirmation of the results of our original study [46] and further description of the sources and characteristics of the artifact. The second experimental design aims to identify the prevalence and associated risks of stimulus artifact occurrence in human ASSR data. The stimulation via speakers is used as a control condition in human experimental design, and the suitability of speakers for stimulation is discussed. Overall, this study will help to create high-quality experimental designs for the neuroscientific ASSR applications with click train stimulation further leading to better interpretability and credibility of the results.

## 2 Methods

The investigation of transduction artifacts necessitates defining potential contributing factors and employing appropriate methods to test their influence on data quality. The factors under investigation and the corresponding methods are summarized in Figure 1.

**Figure 1.**
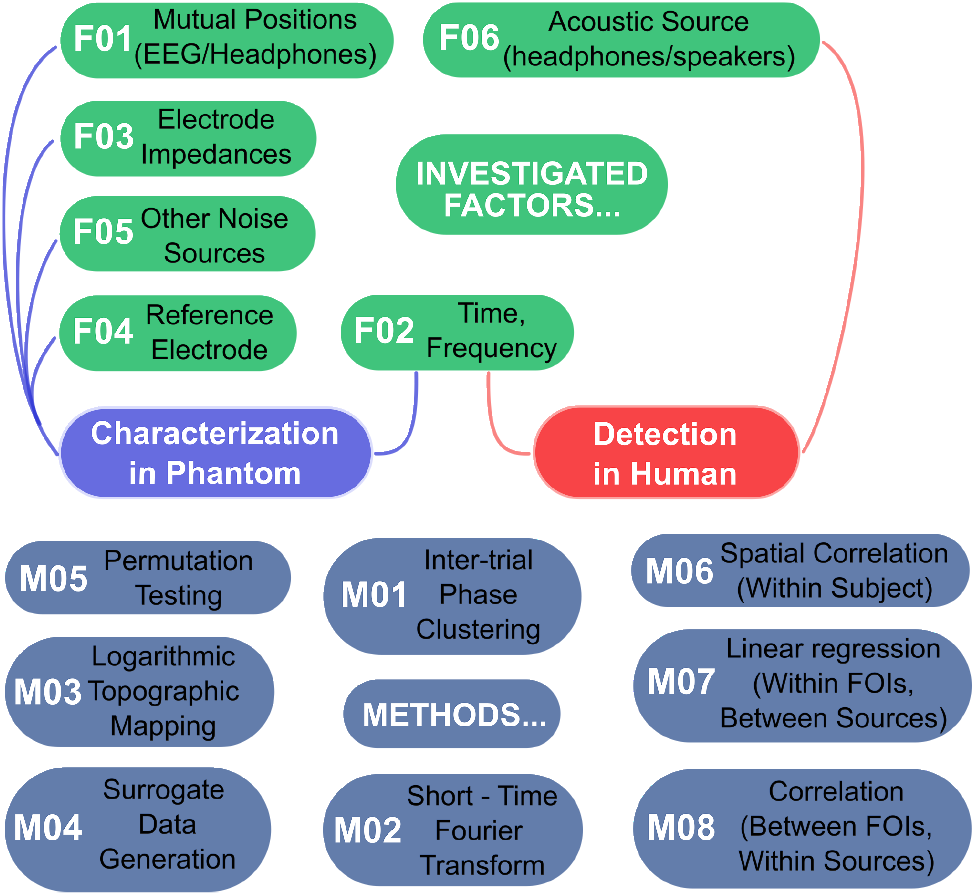
Investigated factors (top) in relation to the artifact characterization in human head phantom and artifact detection in human (middle), and corresponding methods (bottom). Edges in the graph connect factors and methods with corresponding experimental parts: characterization (blue) and detection (red).

In characterizing the artifact, we acknowledge that the noise level is influenced by the relative positions of the EEG and headphone systems (F01). Our experimental designs account for this factor, see section 2.4. The artifact is expected to manifest at specific time intervals (time-locked to the stimulus onset) and specific frequencies (F02). The spectral analysis of the stimulus time series is detailed in section 2.4.1. Additionally, section 2.3 describes the time-frequency analysis using the Short-Time Fourier Transform (M02) and ITPC computation (M01).

The relationship between artifact magnitude and electrode impedance (F03) is examined through topographic mapping (M03) in section 2.4.3. The investigation of different referencing of EEG channels (F04) is elucidated in section 2.2. Lastly, the influence of ITPC values attributed to other sources of noise (F05) was evaluated. For this purpose, surrogate data generation (M04) and permutation testing (M05) are employed, as detailed in section 2.4.2.

Concerning the risks associated with analyzing human ASSR data, with a focus on the stimulus artifact, the primary factor under investigation in experiments involving human subjects is the acoustic source (F06). Spatial correlation analyses (M06) within subjects were conducted, examining differences between spatial patterns of the true brain response and the artifact across different frequencies and acoustic sources.

In the presence of a clean EEG signal, the variance in ITPC across subjects in one acoustic source is expected to be explained by the ITPC in another source. This consideration is incorporated into linear regression analysis within frequencies (M07), see section 2.5.3. Furthermore, the variance in ITPC across subjects at the main frequency of 40 Hz is examined in relation to the variance at higher harmonics. Correlation analysis (M08) between frequencies of interest is employed to assess this relationship, see section 2.5.4.

The initial three sections in methods delineate essential prerequisites implemented in both phantom and human experiments. Specifically, technical equipment, data preprocessing, and the ASSR quantification metric are detailed in sections 2.1, 2.2, and 2.3, respectively.

In accordance with the methods described in section 2.4, a series of systematic approaches was employed to investigate the artifact’s properties. Firstly, the stimulation sound waveform was analyzed to discern its spectral content relevant to the brain response to ASSR stimuli (see section 2.4.1). Subsequently, experiments with a human head phantom were conducted to identify the coupling between the sound source and EEG system, elucidating the artifact’s origin (see section 2.4.2). The spatial properties of the artifact in electrode space were then assessed in relation to the experiment configuration and the distribution of electrode impedance values across the scalp, as outlined in section 3.2.2. Finally, the potential post-hoc correction of noisy EEG records through referencing to the average was examined (see section 2.4.4).

After characterizing the artifact, a series of experiments and data analyses outlined in section 3.2 was conducted to assess the actual risks of data contamination in experiments involving human subjects. Firstly, we ensured the attainment of a standard brain response to ASSR stimuli in both headphones and loudspeaker setups, as discussed in section 2.5.1. Secondly, the spatial properties of brain responses to stimuli presented through headphones and loudspeakers were compared, with the latter considered as the artifact-free reference method (see section 2.5.2). A detailed regression analysis was applied to compare noise levels in EEG data across different harmonic frequencies (see section 2.5.3). Lastly, a between-frequency correlation analysis was conducted to compare noise levels between the two sources of sound stimuli, as described in section 2.5.3.

### 2.1 Technical Equipment

The MagStim EGI GTEN N200 HdEEG amplifier without an electromagnetic shield box was used in our original study [46]. In this study, the signals (testing signals from experiment designs with human head phantom and signals from experiment with human subjects) were acquired with MagStim EGI NetAmp GES 400 HdEEG amplifier with an EGI FICS (Field Isolation Containment System) electromagnetic shield box. We used EGI 256 channel sponge water based net caps. For auditory stimulation, we used an Acoustique Quality M4 DO 2x50 W audio dual-channel amplifier. Acoustique Quality loudspeaker TANGO 85 WHITE speakers and Sennheiser hda 280 headphones were used. The sound volume level was set and measured with an 8922 type GSH 8922 calibrated digital sound meter.

### 2.2 Data Preprocessing

All EEG data preprocessing was performed in the BrainVision Analyzer 2 [47] software for both experimental designs (artifact characterization (see section 2.4), and with human subjects (see section 2.5)).

A standard preprocessiíng pipeline was used for the EEG data from human subjects. Firstly, the signal was filtered by a bandpass IIR filter with default BrainVision Analyzer settings and 1 and 200 Hz cut-off frequencies. A notch filter at 50 Hz was also applied. Next, noisy segments with mainly technical or muscle artifacts were removed. Bad channels were identified and interpolated by spherical spline interpolation with order four and degree 10. EOG (electrooculography) and ECG (electrocardiography) artifacts were then repaired by the ICA method (specifically, a fast ICA algorithm with 35 components). Cleaned EEG data were re-referenced to the average reference and segmented into epochs (−400:850 ms). After preprocessing, EEG data were imported into MATLAB software (MATLAB R2017b) [48] for the next analyses.

Note that the data preprocessing pipeline for the artifact characterization (see section 2.4) was carried out with small differences. Data were filtered only by a highpass filter (no down pass was used) because our interest was also to monitor very high frequencies. Topographic interpolation of bad channels and noisy segment removal was conducted in the same way, but only for technical artifacts. The human head phantom did not create biological artifacts, and no ICA was applied. Data were epoched into segments from -500 to 1000 ms long (related to click train stimuli markers) to analyze the origin of the artifact. The data were not re-referenced, as we also investigated the influence of data re-referencing in our analyses.

### 2.3 ASSR Quantification - Intertrial Phase Coherence

All the analyses applied in this study are based on intertrial phase coherence (ITPC). ITPC, sometimes also called intertrial phase clustering or the phase-locking factor, is a common measure of assessment in neuroscientific ASSR research (see, for example, the meta-analysis in the study [13]). Across experiments, ITPC appears to be more robust than evoked activity [22]. ITPC expresses phase-locking across trials without the influence of the evoked power when ITPC acquires values from 0 (random phases across trials) to 1 (all trials are maximally phase-locked).

The equation used to calculate ITPC is as follows:

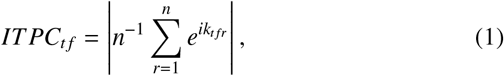

where vertical bars indicate the absolute value, *ITPC*_*t f*_ is intertrial phase coherence calculated in time point *t* and frequency point *f, n* is the total number of trials *r, k* is phase angle, and 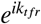 is Euler’s formula for time-frequency point *t f* of trial *r* [49].

Complex Fourier coefficients were used for the ITPC calculation. Specifically, a short-time Fourier analysis was calculated for a time-frequency response of ITPC. The FieldTrip toolbox [50] in MATLAB was used for this analysis with frequencies of interest: 20 : 1 : 250 Hz. The displayed time was counted in the following intervals: -0.25 : 0.05 : 0.74 s (with respect to the onset of the stimulus). The ITPC results calculated from a small number of trials (in contrast to ITPC from a large number of trials) may be affected. Due to data preprocessing (see section 2.2), the number of trials was slightly different among the subjects. It was therefore additionally verified, based on the Rayleigh Z approximation (see the procedure in [51]), that this difference did not affect the ITPC values.

### 2.4 Stimulus Artifact Characterization

In this section, we describe the analyses of the stimulus artifact characterization methodology. Firstly, we investigated the stimulus click train waveform and spectral content to be later identified in the EEG time series and spectrum. Then, the stimulus artifact spatial properties and the influence of the re-referencing to average reference (common ASSR preprocessing step) were investigated.

The phantom experiment designs were targeted to investigate the origin and properties of the stimulus artifact. A wet towel was located between the human head phantom and hdEEG net (see Figure 2) to simulate the impedance of human skin with 95 % of electrode impedance being under 50 kΩ. A similar phantom, specifically the watermelon model, was also used for stimulus artifact identification in the study [30]. In studies [52] and [42], the authors replaced the phantom with severe-to-profound hearing loss participants or participants with occluded ears.

**Figure 2.**
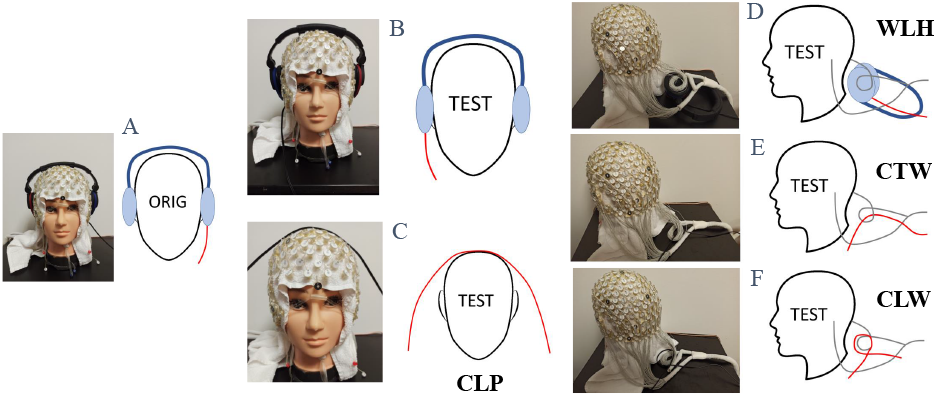
Photo examples and block schemes of the experimental designs used to research the stimulus artifact properties. Specifically for our **original** experiment from the study [46] (A), a new experiment with the same **standard** deployment of the headphones but on the opposite side (B), the headphone cable laying on the human head phantom **CLP** (C), the loop from the hdEEG wires laying on the headphone **WLH** (D), the headphone cable passing through the loop from the hdEEG wires **CTW** (E), and the headphone cable loop lying on the loop from the hdEEG wires **CLW** (F).

#### 2.4.1 Stimulus Waveform Analysis

Firstly, it was necessary to analyze the properties of the used click train stimulation. Similar time-frequency characteristics may be expected for the stimulation artifact. Knowledge of these properties is therefore crucial for subsequent analyses of the stimulation artifact characterization and the risks associated with it for human ASSR experiments. In this section, we used FAM stimulation for comparison with click train stimulation. This is because it is the second most important type of ASSR stimulation, and it is therefore appropriate to compare the properties of both stimulations. However, for the experiments themselves, only click train stimulation, which is used most often in ASSR experiments (see section 1), was used, and at the same time, shows the risk of a stimulation artifact (see section 3.1.1).

The stimulation with a click train stimuli was used in all experiments of this study. The click train stimulus was 500 ms, and it contained 20 noise clicks, each lasting 1.5 ms. The noise clicks were repeated with a frequency of 40 Hz, (see Figure 4 in section 3.1.1). The number of trials was 150 in each condition.

### 2.4.2 Origin of the Stimulus Artifact

Analysis of the origin of the stimulation artifact is possible thanks to special experimental designs using a human head phantom. Therefore, several experiments with the human head phantom were proposed in this study. Firstly, we replicated our original study (see [46]) describing the stimulus artifact: the hdEEG net and headphones were mounted on the human head phantom in the usual way. The EGI GES 400 256 HD EEG system has an extra Field Isolation Containment System, which makes electromagnetic isolation from external electromagnetic noise sources was used. Compared with our original study [46], the headphone cable was placed in reverse, specifically on the right size (see Figure 2B).

The next four tested designs focused on the examination of the artifact emergence. It was expected that the cable or transducer from headphones may generate the stimulus artifact without the influence of another source. Firstly, the design verifying the induction of the artifact from the headphones cable when headphone cable laying on the human head phantom was used, experimental design CLP (see Figure 2C). Secondly, the artifact induction from the headphones transducer to hdEEG was tested by placing the loop from hdEEG wires on the headphone (WLH) (see Figure 2D). Two other experimental designs were intended to verify the induction of the artifact from the headphone cable to the hdEEG wires - the headphone cable passing through the loop from hdEEG wires, CTW (see Figure 2E), and the headphone cable loop lying on the loop from hdEEG wires, CLW (see Figure 2F). We also reanalyzed the data from the original study [46] (see Figure 2A) in order to obtain control results under a large influence of the stimulus artifact on the ASSR.

ITPCs from experimental designs were subjected to classical time-frequency analysis and permutation tests. The permutation tests with a minimum of 400 permutations were calculated to obtain a statistical effect of artifact influence. Specifically, ITPC (see section 2.3) obtained by the FFT from trials with random time-onset (without the influence of click train stimuli) were permuted and compared with phase-locked ITPC (ITPC from trials beginning with the start of click train stimuli, tradition ITPC), see Figure 3. Therefore, we statistically compared phase-locked ITPC (commonly counted ITPC) with ITPCs, representing the noise level of ITPC in the data. Thanks to this analysis, we are able to detect the artifact otherwise invisible to the standard ASSR analysis (time-frequency analysis).

**Figure 3.**
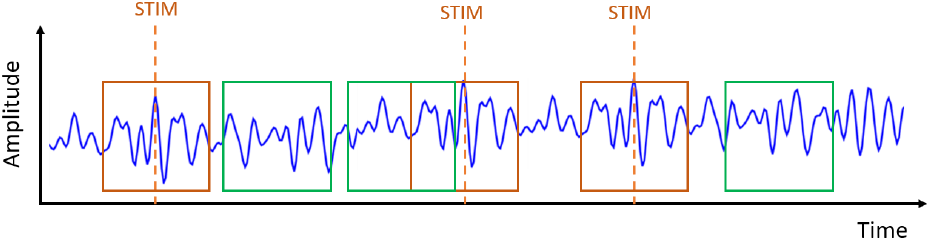
The principle diagram of our permutation test implementation. The blue curve simulates the EEG signal, the orange dash line simulates click train stimuli (STIM), the orange square represents segments phase-locked to stimuli (for computing time-locked ITPC), and the green squares represent random-time obtained segments. Every random permutation utilized a different set of trials with a total number equal to the total number of the phase-locked ones.

**Figure 4.**
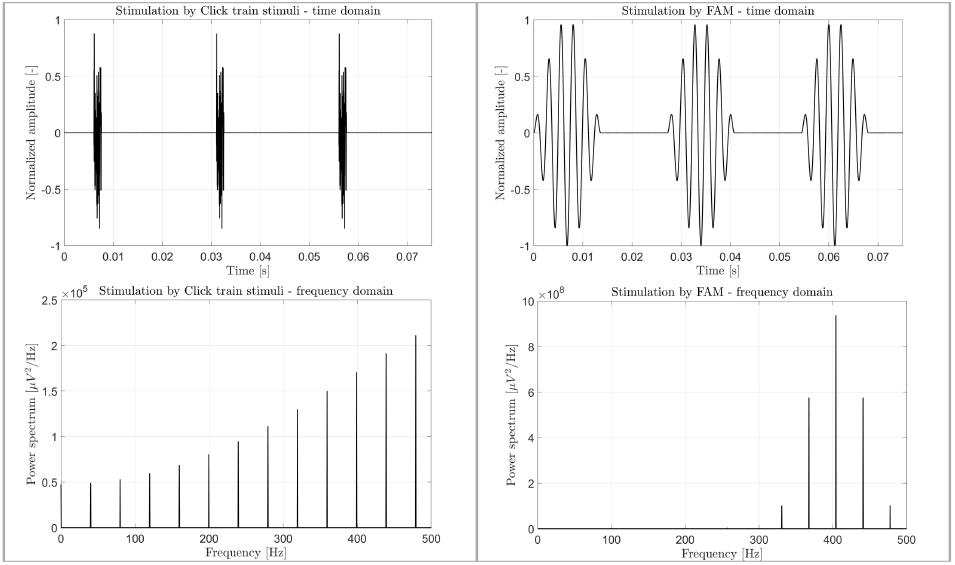
Time characterization of click train stimuli used in this study (top left) and FAM stimuli (top right) displayed for comparison, and power spectral density of click train stimuli (bottom left) and FAM stimuli (bottom right). The power spectral density was displayed for frequencies up to 500 Hz for both ASSR stimuli.

The results of the permutation tests were displayed in two different ways. The histograms of random ITPC (obtained by permutations) were rendered together with ITPC phase-locked to click train stimuli. In Figure 6, different experiment designs for 40 and 120 Hz, frequencies of stimulus artifacts are depicted. We chose 120 Hz rather than 80 Hz because the click train stimulus has higher power at 80 Hz. The trend of the T-statistic is the second way of plotting the permutation test results. The T-statistic (corresponding to p-values lower than 0.05 (statistically significant level)) is plotted for every channel and frequency to see trends obtained from a permutation test across frequencies (the stimulus artifact is manifested only for higher harmonics of 40 Hz).

#### 2.4.3 Stimulus Artifact Spatial Properties

We analyzed the spatial distribution of stimulus artifacts after analysis of artifact sources. The topographic maps were created for each experiment design (see figure 2) reported by stimulus artifacts in previous analyses. Only these experimental designs are able to tell something about the spatial distribution of artifacts. For the same reason, we showed topographic maps only for frequencies reporting stimulus artifacts in these experimental designs. The topographic maps were the caqlculated average across time for the period 0.1 : 0.35 s after the stimulation onset. The ITPC topographic maps were displayed with Robust scaling:

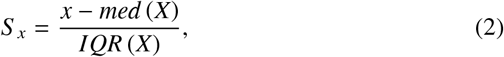

where *x* is an original value from dataset *X, med* (*X*) is the median of dataset *X, IQR* (*X*) is the interquartile range of dataset *X*, and *S* _*x*_ is the final scaling value. Interquartile distance scaling achieved satisfactory results [53] and was mainly used because it is robust to the outliers [54], which occurred in ITPC topographic maps.

The topographic maps of electrode impedance were also displayed. So, we can compare the spatial distribution of the artifact and electrical impedance. The impedance values were plotted after a logarithmic transformation:

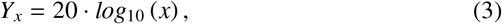

where *x* is an original value, *log*_10_ is a decadic logarithm, and *Y*_*x*_ is a result value after logarithmic transformation.

The logarithmic transformation was applied to compensate for the wide range of impedance values across electrodes.

#### 2.4.4 Effect of Re-Referencing

The analysis of the re-referencing on the average reference is the last part of the human head phantom data investigations. In this part, the influence of the important data preprocessing step was analyzed. We focused on the trend of the T-statistic calculated from the permutation tests (due to the sensitivity to the occurrence of stimulus artifact) of three experimental designs (see section 2.4.2). The original experimental design with the high noise level, testing experimental designs CLP and WLH (see Figure 2), were analyzed because these experimental designs showed stimulus artifact effects in previous analyses. T-statistics were displayed before and after re-referencing to the average reference.

### 2.5 Risks of Artifact Contamination in Humans

The risks of human ASSR experiments (focusing on stimulus artifacts) were investigated after the stimulus artifact characterization analysis. Thereby, we measured human subjects in the ASSR experiment with two acoustic sources (with headphones and speakers as the control condition). Speakers cannot create stimulus artifacts, and we are able to use them as reference methods.

The EEG data of seventeen subjects was collected. Two measured records were removed during data pre-processing for bad quality. The subjects were males and females, between age of 25 and 46 years old. All the subjects were right-handed without any known neuropsychiatric diseases or hearing impairment. The experiment was conducted in accordance with the Helsinki Declaration of Human Rights. The ethical committee of the National Institute of Mental Health (Klecany, Czechia) approved the study. Each subject was, before recording, instructed about the risks and the measurement procedure and signed informed consent. The experiments were performed in an electrically shielded room. Two speakers were placed at the height of the subject’s head. Both speakers were 198 cm away from the subject and 203 cm away from each other.

During both measurements (one with headphones, and one with speakers), the subjects remained with open eyes fixed on a cross. The sound pressure level was calibrated to the loudness of 60 dB SPL (sound pressure level) at the subjects’ ears. Headphones (see section 2.1) were placed on the subject’s head, and click train ASSR stimuli started (see section 2.4.1). The subjects were stimulated only by headphones in this block. The headphones were removed immediately after the end of stimulation. The same stimulation protocol continued in the second block but speakers were used as a stimulus source.

Firstly, we analyzed standard ASSR responses to validate our protocol with headphones and speakers. Subsequently, we analyzed the possibility of detecting the stimulus artifact using spatial analysis. We also investigated the possibility of detecting the artifact by comparison of the noise level during stimulation with different acoustic sources. The character of the detected noise was further analyzed by investigating the harmonic frequency structure.

#### 2.5.1 Standard Response to ASSR Protocol

The standard response to the ASSR protocol was calculated due to the ASSR protocol validation. Measuring the protocol using speakers is one of the reasons for validating the ASSR response.

Two standard analyses were calculated for validation. Firstly, the ITPCs were calculated as a grand mean of all channels and time points from 0.1 to 0.4 s after stimulus for both acoustic sources and all frequencies 40, 80, and 120 Hz. Secondly, time-frequency responses with baseline correction were obtained in the same way as in the experiment for artifact characterization (see section 2.4).

#### 2.5.2 Spatial Properties

Spatial properties were utilized for the analysis of stimulus artifact detection. We created topographic maps in the same way as in the experiment for artifact characterization (see section 2.4.3). The spatial Pearson correlation coefficient was calculated from topographic maps for each subject for frequencies 40 and 120 Hz, separately for each acoustic source. Box-plots were created from these correlation coefficients across subjects (see Figure 10). Additionally, linear similarity (spatial correlation coefficients) between topographic maps for frequencies 40 and 120 Hz in response to both acoustic sources was assessed, and the results are presented in the Supplementary section.

#### 2.5.3 Frequency-Specific Noise Levels

The second part, investigating stimulus artifact detection, used linear regression. This part focused on comparing noise levels for different harmonic frequencies of the 40 Hz stimulus. The artifact occurrence is expected to destroy or decrease the linear dependency between the response evoked by the headphones and speakers across the subjects. The linear regression was calculated to express a dependence of ITPC values from experiments with speakers to ITPC values from experiments with headphones. The Least Square method was used, and the p-value was obtained from the F-test. The null hypothesis of this F-test is that regression coefficients are equal to zero [55] (no relationship between response to stimulation via headphones and speakers).

#### 2.5.4 Acoustic Source-Specific Noise Levels

Linear regression may detect external noise, which corrupts linear dependency between different acoustic sources of stimulation. However, it is important to determine the character of this noise. Then, we are able to identify which artifact led to linear dependency disruption. We investigated the corruption of the harmonic frequency structure (specifically, the structure of 40 Hz and higher harmonic frequencies). The stimulus artifact would be manifested precisely in these structures (differently for stimulation with headphones and speakers, which do not create a stimulus artifact). We calculated cross-frequency correlation coefficients from both acoustic sources. Maps of cross-frequency correlation depict a harmonic frequency structure.

## 3. Results

This section is divided into two parts. Firstly, we present results from measurements on the human head phantom, where we analyzed the stimulus artifact properties, characterize the specific stimulation, the origin of the stimulus artifact, spatial properties of stimulus artifact, and the effect of re-referencing to the stimulus artifact occurrence.

In the second part, we show the results of the ASSR experiment in human subjects investigating the risks in ASSR neuroscientific studies due to the stimulus artifact. The standard analysis of the ASSR is presented for experiment protocol validation. The spatial properties and noise level comparison (due to linear regression) were calculated to detect of stimulus artifacts in human ASSR experiments. Cross-frequency correlation investigated the harmonic frequency structure corruption to identify noise properties (i.e., the noise found by linear regression).

### 3.1 Stimulus Artifact Characterization

#### 3.1.1 Stimulus waveform analysis

It is essential to distinguish the stimulus artifact (generated by the stimuli) from the biological reaction to the ASSR experiment. For this reason, it is important to know the typical properties of the used stimulus, specifically the click train stimulus. In this study, we also described the FAM tone stimulus for comparison. The FAM tone is another stimulation used in ASSR experiments (see, for example, study [56]).

Examples of both stimuli time series are shown in Figure 4, as well as the power spectral densities for both ASSR stimuli. These power spectral density figures focus only on the lower frequencies (frequencies significant for EEG recording). The power spectral density tells us about the frequency characteristic of the stimuli. Thanks to this, it is possible to assume the stimulus artifacts’ frequency characteristics based on specific stimulus types.

#### 3.1.2 Origin of the Stimulus Artifact

The phantom head experiments were designed to reproduce our original study results and show the influence of only headphone cables or transducers, respectively (see section 2.4).

Figures 5 and 6 show the time-frequency response and permutation test results (see section 2.4.2) for three new experimental designs and results from the original experimental design [46]. Two other experimental designs may be found in the Supplementary section for clarity. The results from the permutation test in Figure 5 show a trend of activity being present over more frequency bands for phase-locked ITPC compared to random ITPC. Only T-values representing statistically significant differences from the surrogate distribution were rendered in this figure. We expected stimulus artifacts to be statistically significant only at higher harmonic frequencies of 40 Hz, based on the click train spectral content depicted in Figure 4. Therefore, the figures without any trend in higher harmonic frequencies do not show the stimulus artifact influence and vice versa.

**Figure 5.**
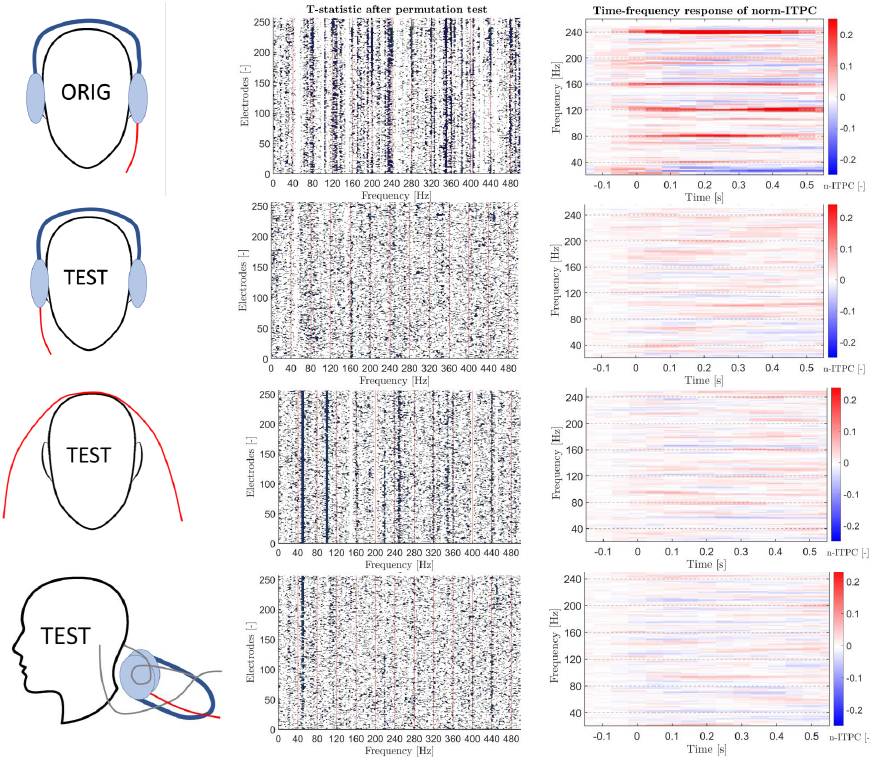
The time-frequency response with baseline correction (right) and T-statistic calculated from the permutation distribution for different frequencies from 0 Hz to 400 Hz and all 256 electrodes (in the middle). Only statistically significant T-values (p-values lower than 0.05) are plotted in the figures of the permutation test. Figures were plotted for experimental designs from our original study, ORIG (top), and new testing designs for this study, TEST. Testing the experimental design with standard EEG settings (second from top), experimental design CLP (third from top), and experimental design WLH (bottom).

Figure 6 also depicts the permutation test results (for the averaged electrodes). This figure shows a difference between phase-locked ITPC and random ITPCs for 40 Hz and 120 Hz, respectively. Therefore, Figure 6 does not show a trend but the results for specific frequencies with the possible occurrence of stimulus artifacts near the EEG frequency range. The quantiles calculated by comparing phase-locked ITPC with all random ITPCs are also plotted in Figure 6.

**Figure 6.**
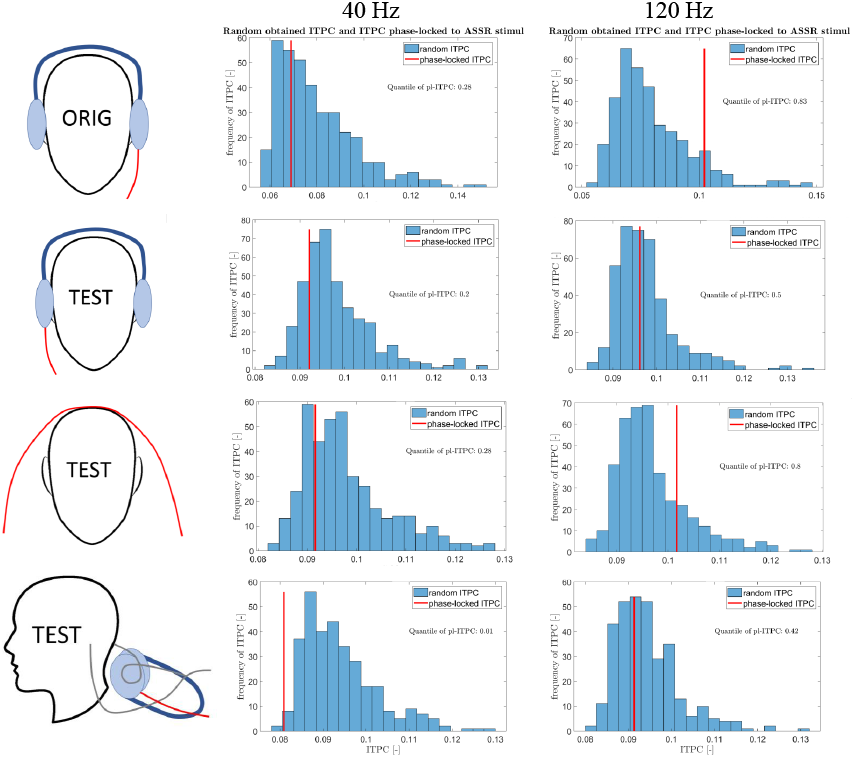
Histogram of ITPC calculated from permuted random segments out of the click train stimuli and ITPC phase-locked to click train stimuli (red line). The graphs were created for frequencies 120 Hz (right) and 40 Hz (in the middle), by averaging across channels. The histograms provide information about the quantile of the ITPC value with respect to the surrogate distribution. Histograms were plotted for both our original design, ORIG (top) and new testing designs for this study, TEST. Specifically, experimental design with standard EEG settings (second from top), experimental design CLP (third from top), and experimental design WLH (bottom).

#### 3.1.3 Stimulus Artifacts Spatial Properties

The spatial distribution of stimulus artifacts was analyzed using topographic maps (see Figure 7). The topographic maps were analyzed only for frequencies potentially affected by stimulus artifacts (see Figure 5). Based on this, the topographic maps were displayed at a frequency of 160 Hz for the experimental design from the original study (see [46]), the experimental design with standard EEG settings, and the experimental design CLP. The results from the next experimental design (experimental design WLH) reported stimulus artifact influence up to a frequency of 440 Hz.

**Figure 7.**
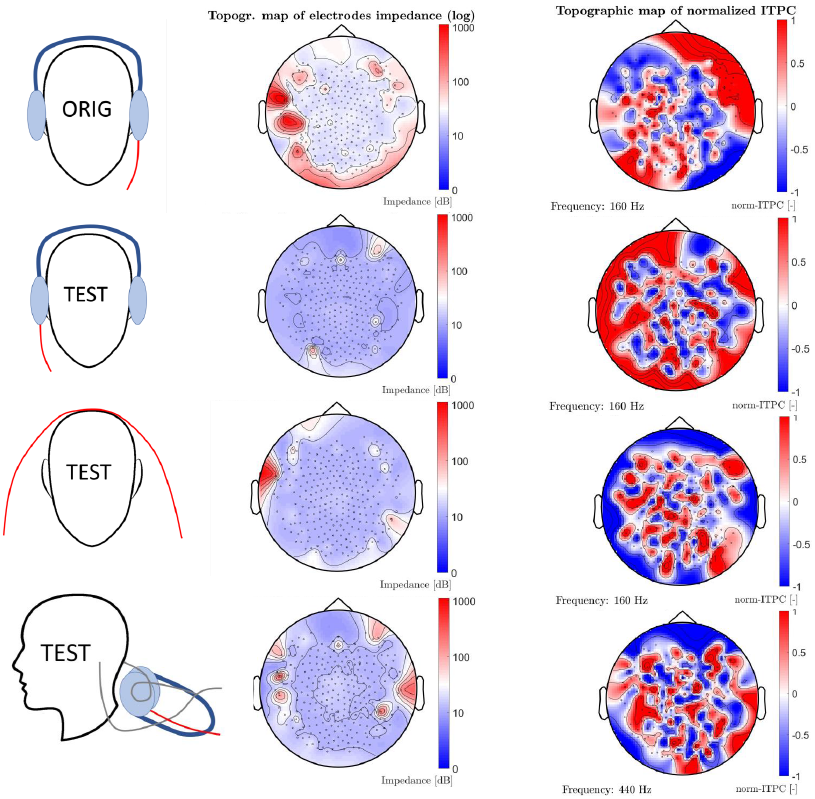
Topographic maps of ITPC at frequencies (160 Hz or 440 Hz) with a significant chance of stimulus artifacts (right) and topographic maps of electrode impedance (in the middle). Impedance topographic maps are plotted after logarithmic transformation. Graphs were plotted for experimental designs from our original study, ORIGIN (top), and new testing designs for this study, TEST. Specifically, the experimental design with standard EEG settings (second from top), experimental design CLP (third from top), and experimental design WLH (bottom).

The topographic maps of electrode impedance were displayed for the same experimental designs. Logarithmic transformation was used for every impedance topographic map. It is possible compare the spatial distribution of stimulus artifacts and electrode impedance by plotting both topographic maps side by side.

#### 3.1.4 Effect of Re-Referencing

Re-referencing to the average of electrodes is a standard preprocessing step in human ASSR experiments. Figure 8 shows the results of the permutation analysis (T-values), same analysis as in section 3.1.2. The results are displayed with and without average re-referencing. Data from three experimental designs were analyzed: a very noisy experimental design from our original study and two less noisy experimental designs with the suspected influence of stimulus artifacts. The testing experimental design with standard EEG settings was omitted in this section due to the low prominence of stimulus artifacts as identified by the permutation test.

**Figure 8.**
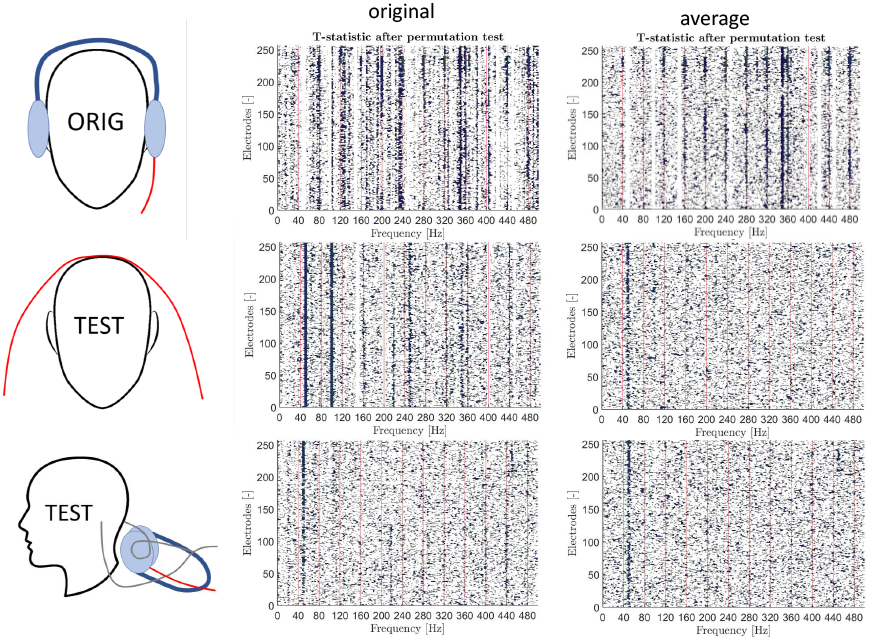
T-statistics calculated from the permutation test for different frequencies from 0 Hz to 400 Hz and all 256 electrodes. The graphs were obtained from records without average re-referencing (in the middle) and with average re-referencing (right). Only statistically significant T-values (p-values lower than 0.05) are plotted in the graphs of the permutation test. Graphs were plotted for experimental designs from our original study, ORIG (top), and new testing designs for this stud, TEST. Specifically, experimental design CLP (in the middle) and experimental design WLH (bottom) are depicted.

### 3.2 Risks of Artifact Contamination in Humans

#### 3.2.1 Standard Response to the ASSR Protocol

The standard time-frequency analysis of the EEG time series was used. Figure 9 shows the grand averaged time-frequency plots of ITPC obtained in the experiment with headphones and speakers. Addition-ally, the box-plots of ITPCs across subjects are depicted.

**Figure 9.**
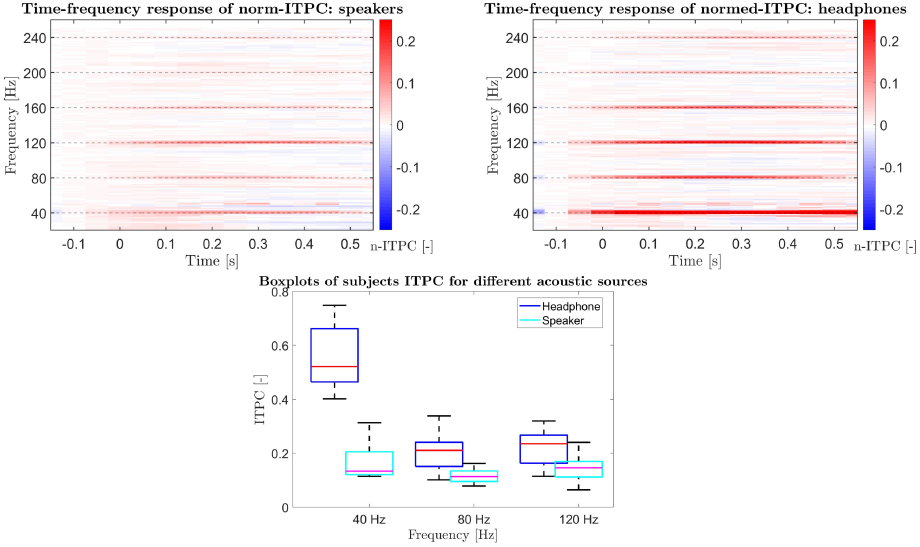
The time-frequency response of ITPC average across subjects for data from the experiment with headphones (top left) and data from the experiment with speakers (top right). The time-frequency response was plotted with baseline correction. The graph below in the middle shows datasets box-plots of ITPC for 40 Hz, 80 Hz, and 120 Hz and experiments with headphones (blue) and experiments with speakers (cyan).

These Box-plots were created for frequencies of interest for the stimulus artifact, specifically at 40 Hz and their higher harmonics (40, 80, and 120 Hz), and for the two acoustic sources (headphones and speakers).

#### 3.2.2 Spatial Properties

Topographic maps of ITPC for 40 and 120 Hz from datasets with different acoustic sources (headphones and speakers) were created, see Figure 10. We selected 120 Hz because it is a relatively higher frequency with respect to the stimulus spectral content (see Figure 4) and is also close to the frequencies of interest.

**Figure 10.**
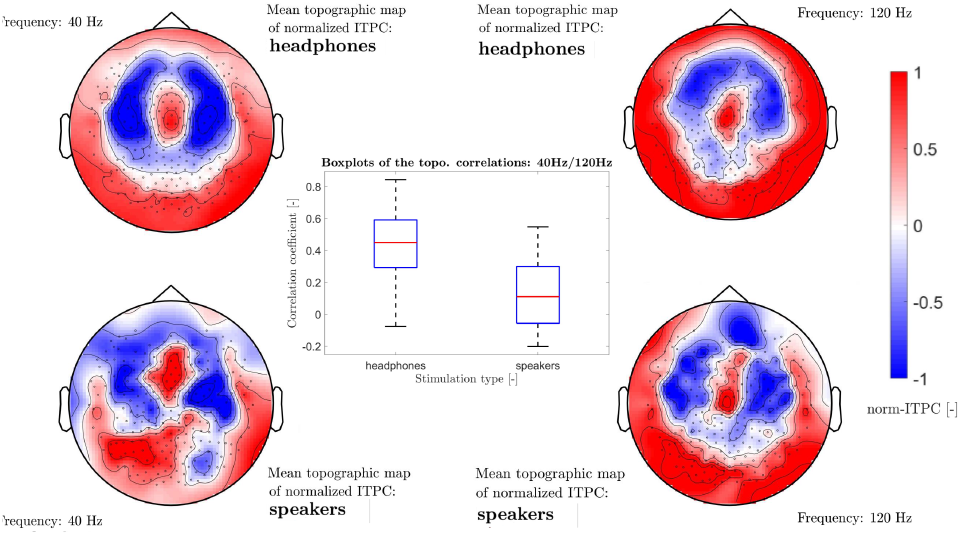
The ITPC topographic maps averaged across subjects for 40 Hz in experiments with headphones (top left), 120 Hz in experiments with headphones (top right), 40 Hz in experiments with speakers (bottom left), and 120 Hz in experiments with speaker (bottom right). Box-plots in the middle depict spatial correlation coefficients between 40 and 120 Hz created from ITPC of 15 subjects for experiments with headphones and speakers across subjects.

The spatial correlation coefficient between the subjects’ topographic maps was calculated. We compared topographic maps between 40 and 120 Hz for the different acoustic sources with the assumption that the correlation coefficient will be lower for experiment with headphones in the case of the influence of the stimulus artifact. Figure 10 shows box-plots of these spatial correlation coefficients for the experiments with headphones or speakers.

#### 3.2.3 Frequency-Specific Noise Level

Linear regression was calculated to compare noise levels at specific frequencies between records with different acoustic sources. Figure 11 shows the linear regression between recordings with different acoustic sources for 40, 80, and 120 Hz (the nearest higher harmonics) and values of the slope of a fit curve and p-values of the F-test. The regression coefficient is not equal to zero at the significance level of 0.05 if the p-value is lower than 0.05, pointing to a relationship between measurements with different acoustic sources.

**Figure 11.**
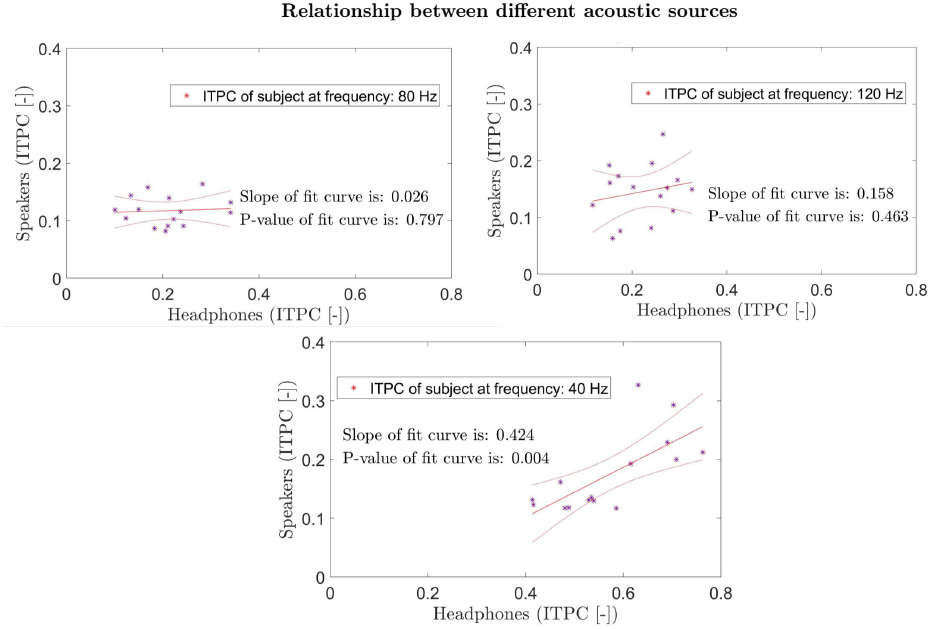
Linear regression of ITPC at frequencies 40 Hz (bottom), 80 Hz (top left), and 120 Hz (top right). Linear regression expresses a dependence of records with speakers to records with headphones (each point in the graph is one subject). In each graph, a value of the slope of the fit curve and the p-value of the F-test is depicted.

#### 3.2.4 Acoustic Source-Specific Noise Levels

The relationship of ITPC between different frequencies average across subjects is displayed in Figure 12, similar to cross-frequency correlation coefficients. This analysis aims to find a spectral character of potential artifact degrading the linear regression of higher harmonic frequencies (see section 3.2.3). Therefore, the cross-frequency correlation is able to display harmonic frequency structure corruption. Figure 12 also includes a cutout of the essential part of the graph (created from recordings during stimulation with headphones).

**Figure 12.**
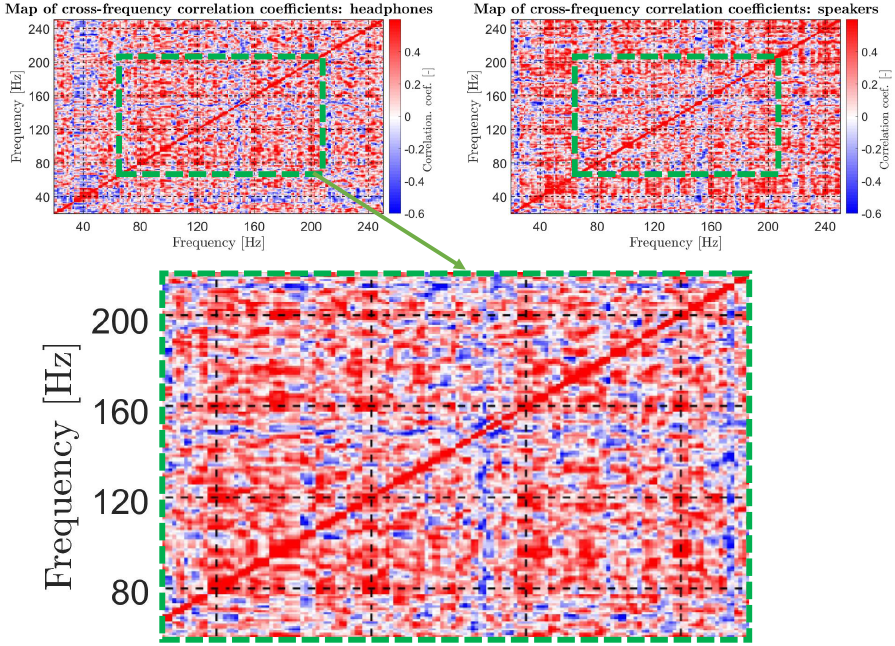
: Maps of cross-frequency Pearson correlation coefficients for records from experiments with headphones (top left) and experiments with speakers (top right). The higher harmonic structure (from 60 to 220 Hz) arising in the case of the headphones is highlighted by green dashed lines and depicted in detail (bottom).

## 4 Discussion

The main aim of this study was to systematically investigate the possible occurrence of stimulus artifact in ASSRs during click-induced stimulation, describing its presence, risks, and properties in neuroscientific ASSR experiments. In our original study [46], we demonstrated the existence of an artifact in a pilot experiment on a phantom of the human head. Based on these results, this study mainly focused on characterizing the stimulus transduction artifact (stimulus artifact) occurring in response to click stimulation on an extended set of measurements with phantoms and evaluating possible outcomes in experiments with human subjects. This study was divided into two parts. The first part aimed to investigate stimulus artifact properties essential to possible artifact avoidance. The second part aimed to investigate the artifact in neuroscientific ASSR data from experiments on human subjects.

### 4.1 Stimulus Artifact Characterization

In this experimental part of the study, we investigated the stimulus artifact characteristics during ASSR using a click train.

Firstly, we investigated the responses to click train stimuli as they are well-tested and frequently applied in neuroscientific research. As shown in Figure 4, the modulating envelope of the FAM was 40 Hz, but the carrier frequency was near 400 Hz. The FAM power spectrum thereby interfered with much higher frequencies than the EEG signal. Compared to that, the whole spectrum of the click train stimuli was created from the peaks with a frequency distance of 40 Hz (see Figure 4). Therefore, the click train stimuli peaks also occurred at a frequency of 40 Hz and its higher harmonics frequencies, and could therefore produce the stimulus artifact with similar properties to the brain reaction as investigated with ASSRs, causing spurious responses to click train stimuli measured by EEG. However, it should be noted that the overall power spectrum of a particular click train stimuli depends on the character of white noise.

Furthermore, we suggested five new experimental designs and compared them (see section 2.4) to our original experimental design. The original experimental design from our study [46] included measurements of hdEEG using a system without electromagnetic isolation and showed a very significant influence of the stimulus artifact. Two testing experimental designs (experimental design CTW and experimental design CLW) did not show the effect of stimulus artifact, suggesting that inductive or capacitive coupling between EEG harness and headphone cable are not the main sources of the transduction artifact. The following three testing experimental designs represent standard EEG settings (headphones on the head), allowing the test of the influence of only the headphone cable (experimental design CLP) and the influence of the headphone transducer (experimental design WLH).

As seen in Figure 5, we measured the signal with the stimulus artifacts in our original experiment. This is reflected in significantly high ITPC at higher harmonics of 40 Hz in the time-frequency response. This effect may be due to only artifacts bound to the click train stimulation because of the non-existent physiological signal in the human head phantom. On the other hand, none of the other testing experimental designs showed signs of stimulus artifact presence in the time-frequency response (see Figure 5). For this reason, we also focused on deeper statistical analysis the occurrence of stimulus artifacts. Figure 5 shows T-values derived from the permutation test with statistical significance across frequencies and channels, allowing observation of the structure of phase-locked ITPC across frequencies and electrodes. The results of our original experiment are highly noisy, with a prominent influence on higher harmonic frequencies of 40 and 50 Hz. The experimental design CLP has a distinctive pattern for higher harmonics of 40 Hz. The exception is primarily in higher harmonics common to 40 and 50 Hz, where the result is probably influenced by the 40 Hz artifact and line noise combinations. The experimental design WLH also has a 40 Hz higher harmonics pattern in permutation analysis, but only at very high frequencies. The testing experimental design with standard EEG settings did not show the presence of stimulus artifacts, as seen in Figure 5.

Importantly, the noise ITPC distribution was comparable across all records (see Figure 6), and higher (though not significant) quantiles of phase-locked ITPC for higher frequencies could be observed. This aligns with the permutation test from Figure 5, pointing to the fact that the stimulus artifact appears prominently at higher frequencies (for our click train stimulus implementation).

The stimulus artifact was not observed in the standard EEG settings, except at an isolated frequency peak of 160 Hz. However, two out of five testing experimental designs (experimental design CLP and experimental design WLH) measured on the EEG system with electromagnetic isolation exhibited the presence of the artifact (see Figure 5). The artifact was detected by a statistical evaluation while not being visible in the standard time-frequency analysis, pointing to the danger of interpreting the artifact as a brain response. The source of stimulus artifact may be both in the cable and transducer of the headphones; however, it is surprising that the headphone cable contributes to the generation of stimulus artifact more than the influence of the headphone transducers that had a mild effect in our analysis.

Furthermore, we analyzed the spatial properties of stimulus artifacts after verification of the artifact’s existence. We focused on the frequencies with a high risk of stimulus artifact from the permutation test (see Figure 5). The frequency of 160 Hz was selected for three experimental designs (our original study, testing design with standard EEG settings, and testing experimental design CLP). The frequency of 440 Hz was selected for the experimental design WLH because for 160 Hz there was no noticeable influence of the artifact. Here, the interesting information is the influence of the high electrical impedance of electrodes on the stimulus artifacts’ spatial properties. So, we also created topographic maps of electrode impedances.

Figure 7 shows the individual topographic maps. Neither the experimental design with a cable laid on a human head phantom nor the experimental design of a loop of hdEEG wires laid on a headphone transducer resulted in the expected spatial distribution. The headphone cable was on the right side of the human head phantom in standard EEG settings from our original study and on the left side in standard EEG settings from our actual testing experimental design. The ITPC had higher amplitude on the same sides of topographic maps for both experimental designs. This may indicate the influence of the headphone cable on the stimulus artifact origin. Nevertheless, it is necessary to pay attention to non-existing trends in the permutation analysis of the testing experimental design with standard EEG settings. It is clear that some artifacts have an effect here at a frequency of 160 Hz, but we do not know if it is a stimulus artifact. So, the relationship between spatial distribution and headphone cable may be only by chance. A non-existing relationship between the spatial distribution of the stimulus artifacts and the electrode impedance may be seen in Figure 7.

Re-referencing the EEG signal to the average reference is a standard part of ASSR neuroscientific data preprocessing. For this reason, we analyzed the influence of re-referencing to the stimulus artifact (see Figure 8). The T-statistic calculated from permutation tests was utilized due to its sensitivity to stimulus artifacts. The stimulus artifact effect was not proven in the testing experiment with standard EEG settings. Therefore, we did not present the re-referencing effect to these experimental designs in the main text but in the Supplementary. The disappearing pattern of stimulus artifacts for testing experimental designs is evident in Figure 8. This suggests that re-referencing to an average reference may suppress stimulus artifacts in lightly noisy records. However, the re-referencing to the av-erage reference emphasized the stimulus artifact in the highly noisy record from our original experimental design. This is because of the significant effect of stimulus artifacts only in small spatial regions. The average reference then projects the stimulus artifact to other channels.

To conclude this section, we discovered the permutation test is more sensitive to the stimulus artifact than to the time-frequency response. In some experimental designs using EEG with electromagnetic isolation, stimulus artifacts occurred in permutation tests but not in time-frequency response. Some of the experimental designs with the same settings did not prove the stimulus artifact. We cannot determine, at the moment, why that is, which is a potential risk for ASSR experiments.

Our results indicate that the headphone cable is a significant source of stimulus artifacts. However, headphone transducers also have a measurable influence on the origin of the stimulus artifact. Therefore, using the earphones with a plastic tube is insufficient to suppress the stimulus artifacts. We did not prove the influence of electrode impedance on the stimulus artifact in this study. We show that the average re-referencing is able to suppress stimulus artifacts. On the other hand, average re-referencing may carry the stimulus artifact into all electrodes if the stimulus artifact is expressive and located in a concrete spatial area. Therefore, it is essential to watch for the spatial character of the artifact because there is a risk of false positive results.

### 4.2 Risks of Artifact Contamination in Humans

The main aim of this study was to investigate the potential confounds of stimulus artifacts on ASSR responses in humans. After confirming the possible occurrence of stimulus artifacts in ASSR experiments on phantoms, we analyzed EEG data from an experiment with human subjects. The problem here was the similarity between the frequency pattern of the stimulus artifact and the frequency character of the brain response. For this reason, the distinction between stimulus artifact and the ASSR response is complicated, and we had to proceed with a series of original analyses. We performed both headphone and speaker stimulation (reference recordings) for this reason. The average re-reference was used to comply with the typical ASSR experiment conditions.

Firstly, we validated our experiment design. We display average time-frequency responses for both acoustic sources. A similar time-frequency response as in other studies (for example, [12, 22]) may be seen in Figure 9 for the experiment with headphones. The ASSR response was shown for data obtained during stimulation with speakers as well, even for higher harmonic frequencies of 40 Hz (see Figure 9). However, the signal-to-noise ratio for 40 Hz is much lower than the headphones. Multipath propagation may be the reason for this. Therefore, in ASSR experiments with speakers, it is essential to emphasize the details of experimental design and the acoustic properties of the recording room. The box-plot in Figure 9 shows a low signal-to-noise ratio for higher harmonic frequencies in both acoustic sources. The human brain response to ASSR stimulation may arise at higher harmonic frequencies (the proof is the occurrence in experiments with speakers) where stimulus artifact may have profoundly influence the signal.

The topographic maps at a frequency of 40 Hz in Figure 10 also confirm the typical topographic distribution of ASSR response for both acoustic sources. The topographic maps at 120 Hz (a higher harmonic frequency of 40 Hz near the edge of a typical EEG frequency band) had the same spatial characteristic. The high ITPC in the frontocentral area was evident in all topographic maps (same as in, for example, in [23, 57]). The pattern of the neural ASSR response is proven in the topographic maps, but there was no noticeable effect of the artifact.

For this reason, we analyzed the more profound influence of stimulus artifacts on spatial distribution. We based this analysis on four premises. Firstly, the stimulus artifact does not have spatial characteristics typical for human ASSR (see section 3.1). Secondly, the stimulus artifact for our click train stimuli has a higher influence in higher harmonic frequencies than in 40 Hz (see section 3.1). Thirdly, the speaker does not create a stimulus artifact, but the stimulation induces ASSRs (see section 3.2.1). Finally, the experiments with both acoustic sources immediately followed each other without any other setting changes. Therefore, if stimulus artifacts influence the spatial distribution of the response to stimulation with headphones, we assumed a higher spatial correlation between 40 Hz and 120 Hz for experiments with speakers to occur. The results of spatial correlation across subjects are depicted in the boxplot in Figure 10. It is possible to see that the spatial correlation is opposite to our expectation. We also created box-plots of the spatial correlation between speakers and headphones for 40 Hz and 120 Hz (a figure is included in the Supplementary) and expected the higher spatial correlation to be observed at 40 Hz. That was not the case. Hence, our experiment did not confirm the spatial influence of stimulus artifacts. The stimulus artifact is not present in this dataset, or, alternatively, the stimulus artifact spatial influence was suppressed by the low signal-to-noise ratio of ITPC in an experiment with speakers, possibly amplified by average re-referencing. We further analyzed this pattern expecting that stimulus artifacts would destroy the relationship between ITPC measured from experiments with different acoustic sources at higher harmonic frequencies (affected response to headphone stimulation only), but would not destroy this relationship for a frequency of 40 Hz. Figure 11 shows the linear regression of individual ITPC between experiments with different acoustic sources for 40 Hz, 80 Hz, and 120 Hz frequencies, confirming our expectations (the Supplementary includes confirmation of our expectation also for frequencies 160 Hz and 240 Hz, 200 Hz is influenced by a line noise artifact). Regression analysis results indicate the influence of some noise, which suppresses the ITPC relationship as we assumed in the occurrence of a stimulus artifact. The lower signal-to-noise ratio at higher harmonic frequencies may also explain this phenomenon. Future work with a larger number of subjects will bring certainty about the cause of the regression analysis results.

We also analyzed the cross-frequency correlation coefficient in records from experiments with both acoustic sources. Compared to the regression analysis, we looked at the link between frequencies for different acoustic sources separately (see Figure 11). In the case of stimulus artifact influence, we assumed an existing pattern between higher harmonic frequencies of 40 Hz in records from an experiment with headphones but a non-existing (same) pattern in an experiment with speakers. This expectation was confirmed, as may be seen in Figure 12. In the experiment with speakers, the 40 Hz frequency was correlated with higher harmonic frequencies, but there was no strong correlation of higher harmonic between themselves. On the other hand, in the experiment with headphones, there was a correlation between higher harmonic frequencies, but not between the frequency of 40 Hz and its higher harmonics. These results point to a hidden noise pattern in higher harmonic frequencies of 40 Hz in experiments with headphones. Therefore, it confirms the presence of the stimulus artifact in the human dataset despite the fact of the average re-referencing.

ASSR experiments utilizing speakers do not create stimulus artifacts, and we confirmed that they may evoke human ASSR. However, it is crucial to focus on the correct settings of experiments with speakers to get a higher signal-to-noise ratio. The stimulus artifact was not observed in the spatial characteristics of the human ASSR data after average re-referencing. Despite that, an artifact structure was observed at higher harmonic frequencies in the correlation analysis. Fortunately, the influence was not found at the most standard frequency of 40 Hz. Therefore, signals at the higher harmonics, if analyzed in any ASSR experiment (for example, as in the study [22]), have to be treated with special care to prevent the occurrence of stimulus artifacts. The headphone cables should be routed as far as possible from the EEG wires, and headphone transducers should be placed as far as possible from the EEG electrodes. A better result may be achieved by using waveguides, which lead, on the other hand, to waveform distortion due to their limited bandwidth. Once the EEG signal is contaminated by the stimulus artifact, it is very difficult to clean it due to its similarity to both amplitude and phase spectral content. At least, a within-subject permutation test similar to our analysis (see Figure 3) is recommended to rule out the presence of the artifact.

## 5 Conclusion

This study was aimed at the analysis of stimulus transduction artifacts (stimulus artifacts) and their influence on human ASSR data. We analyzed ASSRs in response to click train stimuli at 40 Hz. We found significant stimulus artifacts in our previous original study with the human head phantom [46]. In this paper, we first confirmed, extended, and generalized our original results and then analyzed the properties of stimulus artifacts in experiments with the human head phantoms. In the end, we focused on the possible confounds during ASSR experiments with human subjects.

The stimulus artifact looked insignificant when using an EEG device with electromagnetic isolation and could be suppressed by average re-referencing (a common preprocessing step in ASSR analysis). However, average re-referencing may highlight the stimulus artifact across electrodes in a situation of a significant stimulus artifact in a concentrated spatial area. The source of the stimulus artifact was identified in the headphone cable, but some noise similar to the stimulus artifact was also generated from headphone transducers. Therefore, using headphones without cables may not be enough for suppressing the stimulus artifact. Experiments with human subjects did not prove the stimulus artifact at a frequency of 40 Hz. On the other hand, stimulus artifacts may be hidden in higher harmonic frequencies, and the induction of stimulus artifacts was variable in experiments with the human head phantoms.

While following the basic rules of the experiment, the classical analysis of ITPC at 40 Hz should not be affected by stimulus artifacts. We recommend placing the headphone cable as far as possible from EEG electrodes and checking the topographic maps of the response at a higher harmonic frequency of 40 Hz (for example, 120 Hz) before average re-referencing. We also proved that stimulation with speakers may create ASSR responses. However, the signal-to-noise ratio is susceptible to the experimental design. Caution is needed to find the correlation effect in data and mainly analyze higher harmonic frequencies of 40 Hz. The high ITPC outside the frontocentral area may alert to significant stimulus artifacts in the data. The within subject permutation test and cross-frequency correlation analysis may also help. These accessible analyses may prevent false positive results after average re-referencing.

## Supporting information

Supplementary Figure 3 represents box-plots of the spatial correlation coefficient.

Supplementary Figure 1 represents the time-frequency analysis and T-statistic for experimental designs CTW and CLW.

Supplementary Figure 2 represents the time-frequency analysis and T-statistic after data re-referencing.

Supplementary Figure 4 represents the linear regression of ITPC at frequencies 160 Hz and 240 Hz.

## CRediT authorship contribution statement

**Jan Strobl:** Formal analysis, Programming, Methodology, Writing – methodology, discussion. **Vojtech Viktorin:** Conceptualization, Writing – introduction. **Marek Piorecky:** Formal analysis, Writing – results. **Inga Griskova-Bulanova:** Supervision, Writing – conclusion. **Jan Hubeny:** Methodology, Data Curation. **Martin Brunovsky:** Validation, Funding acquisition. **Tomas Palenicek:** Project administration. **Vlastimil Koudelka:** Conceptualization, Supervision, Programming.

## Declaration of competing interest

The authors declare that they have no known competing financial interests or personal relationships that could have appeared to influence the work reported in this paper.

## Data available

Data will be available upon request.

## Code available

The code is available at the following link: https://github.com/strobja/STart_ASSRexp. When using it, please follow the license terms and cite the code according to the information in the link.

## Acknowledgments

This project was supported by the Ministry of Health Czech Republic -DRO 2021 (National Institute of Mental Health - NIMH, IN: 00023752), the Grant Agency of the Czech Technical University in Prague, reg. no. SGS22/200/OHK4/3T/17 and SGS24/110/OHK4/2T/17, the Czech Science Foundation project No. 21-32608S. The publication was also supported by Specific University Research, Czech Ministry of Education, Youth and Sports (project 260648/SVV/2024). The publication was also supported from ERDF-Project Brain dynamics, No. CZ.02.01.01/00/22_008/0004643.

## Notes

https://github.com/strobja/STart_ASSRexp

### Competing Interest Statement

The authors have declared no competing interest.

