## Supplementary figures and images for "Unveiling Stimulus Transduction Artifacts in Auditory Steady-State Response Experiments: Characterization, Risks, and Mitigation Strategies"

### Supplementary Figure 1 represents the time-frequency analysis and T-statistic for experimental designs CTW and CLW.

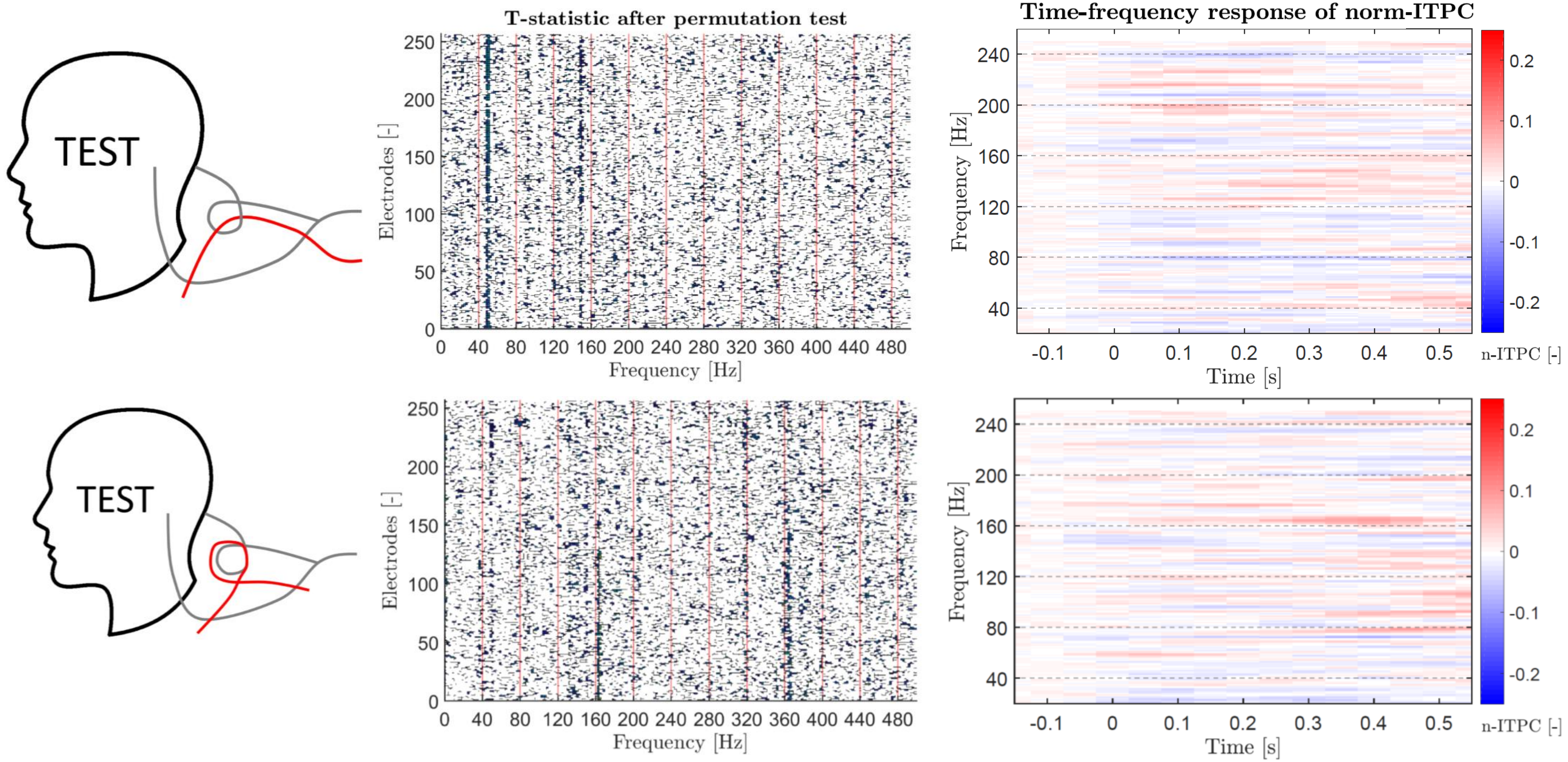

### Supplementary Figure 2 represents the time-frequency analysis and T-statistic after data re-referencing.

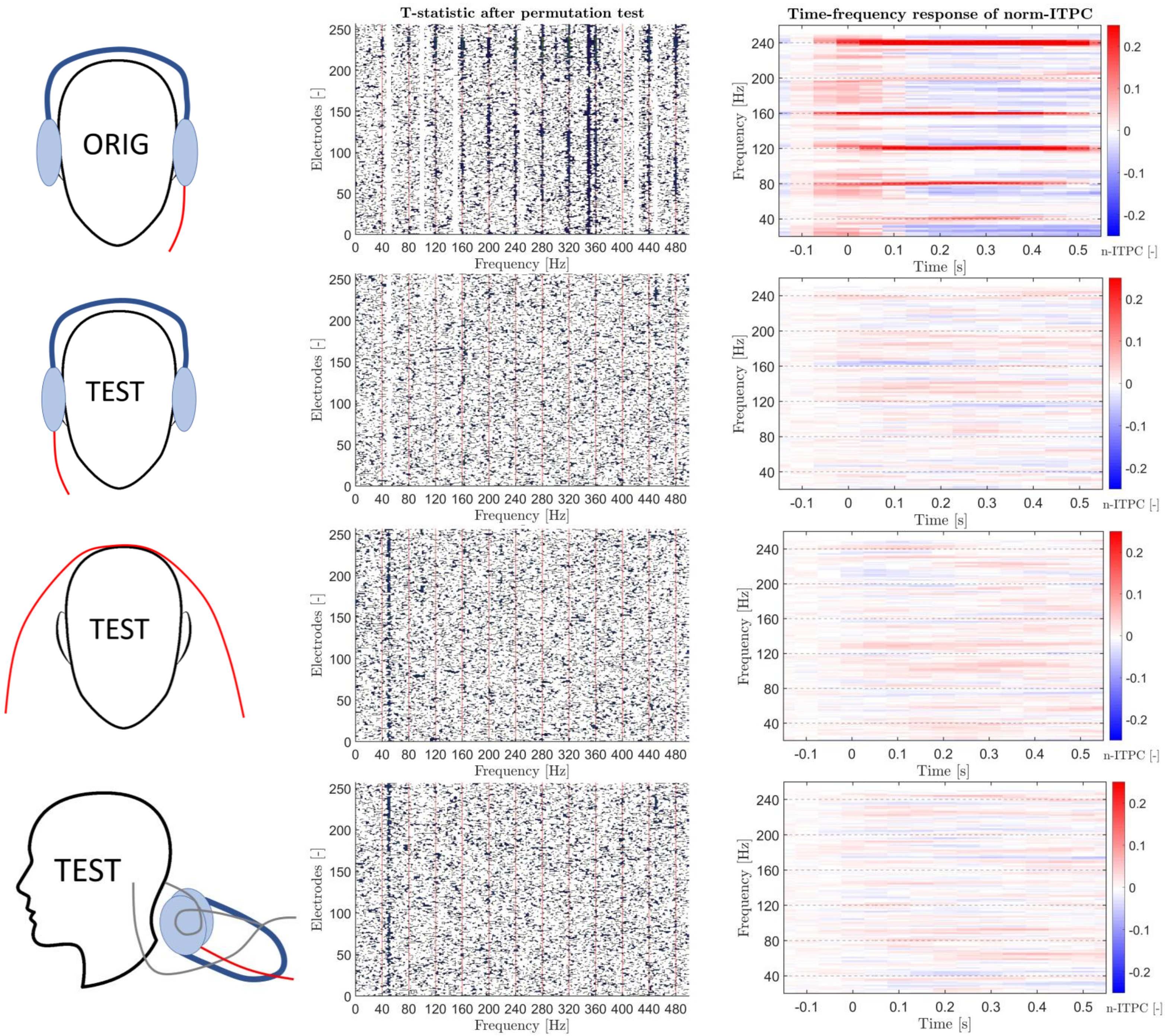

### Supplementary Figure 3 represents box-plots of the spatial correlation coefficient.

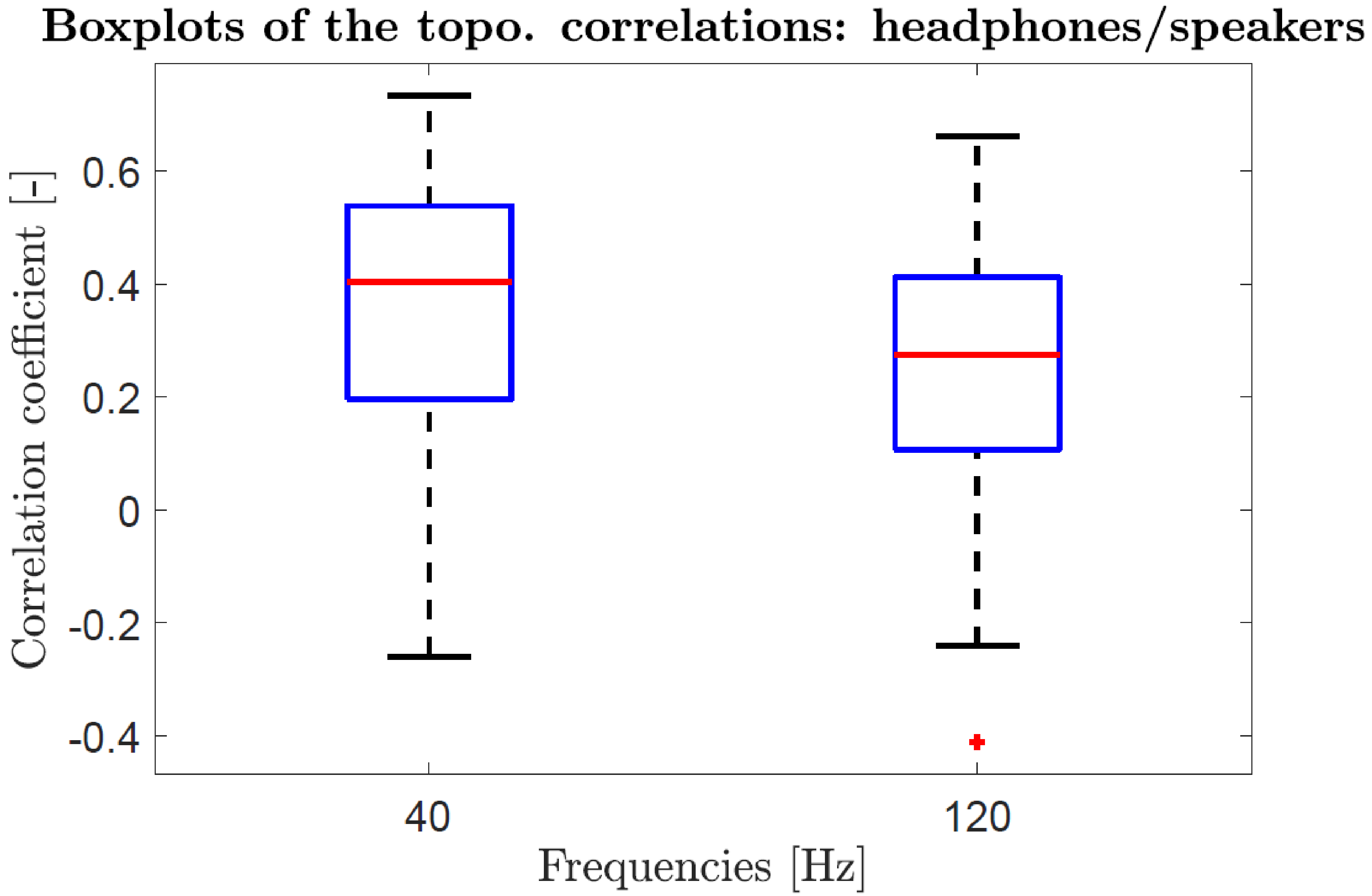

### Supplementary Figure 4 represents the linear regression of ITPC at frequencies 160 Hz and 240 Hz.

#### Relationship between different acoustic sources

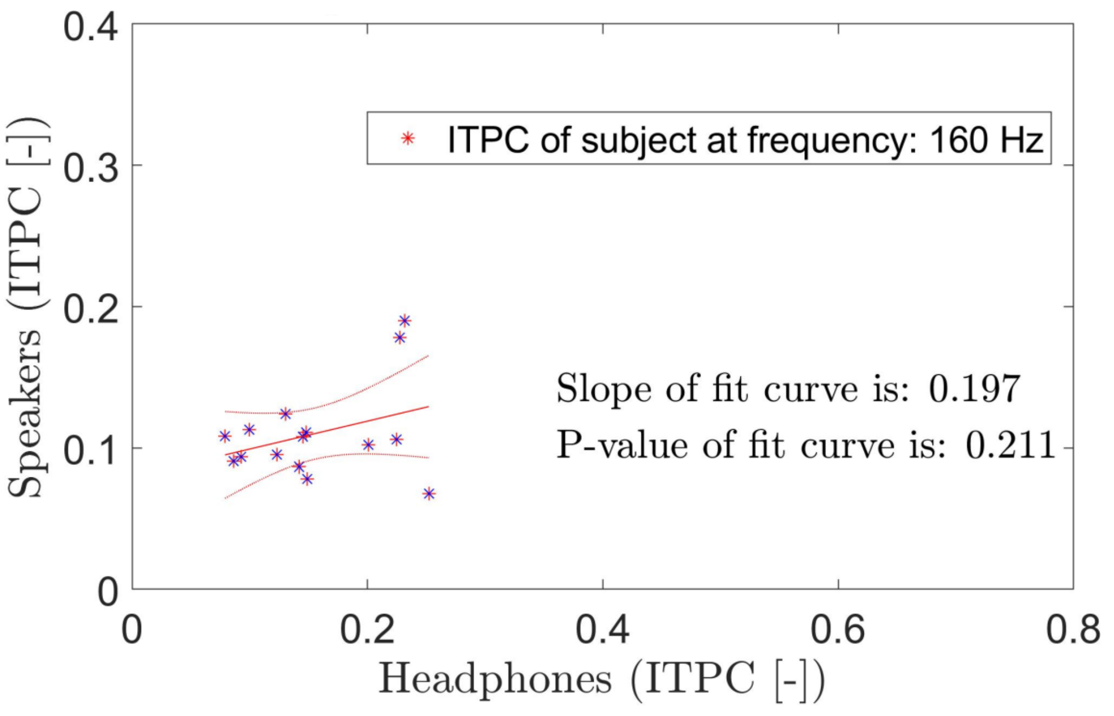


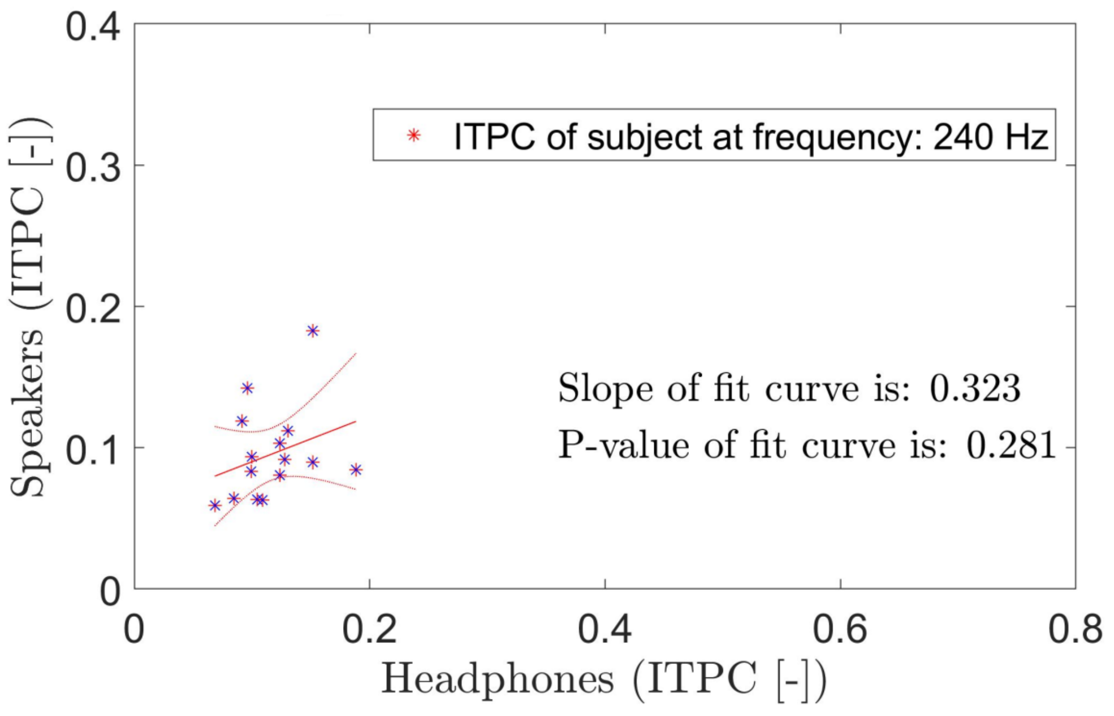
